# Male evolution under relaxed selection: Evidence for degeneration in sperm produced by male snails from asexual lineages

**DOI:** 10.1101/556357

**Authors:** Joseph Jalinsky, John M. Logsdon, Maurine Neiman

## Abstract

How drastic changes in selective regimes affect trait evolution is an important open biological question. We take advantage of naturally occurring and repeated transitions from sexual to asexual reproduction in a New Zealand freshwater snail species to address how relaxed selection on male-specific traits influences sperm morphology. The occasional production of male offspring by the otherwise all-female asexual lineages allows a unique and powerful opportunity to assess the fate of sperm traits in a context where males are unnecessary. These comparisons revealed that the sperm produced by “asexual” males are markedly distinct from sexual counterparts. In particular, the asexual male sperm harbored markedly higher phenotypic variation and was much more likely to be morphologically abnormal. Together, these data suggest that transitions to asexual reproduction might be irreversible at least in part because male function is likely to be compromised. More broadly, our results are consistent with a scenario where relaxed selection translates into rapid trait degeneration.

---

Evolution by natural selection generates adaptation and often underlies the initial appearance and subsequent elaboration of complex traits. While Charles Darwin was one of the very first to link selection and adaptation, he also recognized the importance of variation away from presumed optimal phenotypes (Darwin 1859). A striking example of this phenomenon is the eye of *Astyanax mexicanus*, a freshwater fish characterized by both surface and cave ecotypes. One of the main differences between the *A. mexicanus* surface and cave forms is that the cave dwellers have lost eye structures, likely linked to the perpetual absence of light in their habitat (Jeffery 2005, Cartwright *et al.* 2017). The process that yields the dismantling of once useful traits, so-called regressive (or degenerative) evolution, is generally thought to be a consequence of a shift in selection regime and can result in the degeneration of ancestrally important traits. Examples of trait loss *via* regressive evolution are common in nature and across diverse taxonomic groups, including humans (Prout 1964, Wilson 1982, Remko *et al.* 2005, Ansaloni & Catena 2009; Stanley 2014). Transitions to relaxed selection are often attributed to environmental or life-history alterations that render historically important phenotypes irrelevant (Lahti *et al.* 2009; Ellers *et al.* 2012; Helliwell 2013).

We investigate how shifts in reproductive mode (here, sexual to asexual reproduction) alter the selective regime experienced by traits that are used only for sexual reproduction. Our particular focus is on the nature and extent of the loss of “maleness” in asexual lineages where females reproduce without any need for males. In this scenario, males, and the genes underlying male-specific traits, are presumably extraneous (Carson *et al.* 1982). This is the basis for the expectation that in these asexual lineages, traits needed only for sexual reproduction (“sex-specific traits”; *e.g.*, genes used only for meiosis or male-specific traits like sperm) (Schurko & Logsdon 2008; Tvedte *et al.* 2017) will experience a shift from purifying to relaxed selection at the onset of asexuality. If maintenance of these now-neutral sex-specific traits is not otherwise costly, and if the genes underlying the sex-specific traits are not pleiotropic with respect to traits important outside the context of sex, the sex-specific traits will subsequently evolve primarily via genetic drift and mutation instead of selection (van der Kool & Schwander 2014). If maintenance of these traits does involve cost, selection favoring trait loss in the asexual lineage can contribute to the rate of trait degeneration. These expectations have been evaluated with respect to sexual signaling in bush crickets (Lehmann *et al.* 2007) and wasps (Pannebakker *et al.* 2005), sperm-storage organs and fertilization success of males produced by otherwise all-female asexual lineages in stick insects (Schwander *et al.* 2013), and sperm formation in oribatid mites (Taberly 1988; Maraun 2003; Heethoff 2007). While these studies indicate that transitions to asexuality often but not always (*e.g.* Lehmann *et al.* 2007) underlie sex-specific trait degeneration, the conclusions that can be drawn are limited by the inability to directly compare the sexual and asexual forms (*e.g.*, sexuals and asexuals are allopatric, are different species, or the asexuals are hybrids).

How sperm and seminal fluid proteins evolve under relaxed selection is of particularly broad interest because these traits are among the most important mediators of fertilization, sperm competition, sexual conflict, and speciation (Swanson & Vacquier 2002). Accordingly, characterizing sperm evolution in the context of asexuality will provide a rare glimpse into the consequences of rapid shifts in selective regime for traits that are usually critically important determinants of evolutionary trajectories.

There are two major barriers facing the characterization of the evolution of sperm and related traits in an asexual background. First, obligately asexual lineages derived from dioecious taxa are typically all or nearly all females (*e.g.*, Gottlieb & Zchori-Fein 2001; Schon *et al.* 2009; Neiman *et al.* 2012). Second, even if male asexuals exist, powerful and rigorous characterization of how relaxed selection influences sperm evolution requires the direct comparisons that can only come from sperm produced by ecologically and phenotypically similar males from closely related sexual lineages. Perhaps because many asexual taxa are either of hybrid origin or do not have close sexual relatives (reviewed in Neiman & Schwander 2011), this type of study has yet to be conducted.

We are able to overcome both of these barriers by taking advantage of the occasional production of male offspring by obligately asexual *Potamopyrgus antipodarum* (Neiman *et al.* 2012). This New Zealand freshwater snail provides a powerful means of directly evaluating the impact of shifts in reproductive mode because phenotypically and ecologically similar obligately sexual and obligately asexual snails regularly coexist in natural populations (Lively 1987; Neiman *et al.* 2011), enabling direct comparisons between sexual and asexual individuals, genomes, and populations. Asexual *P. antipodarum* are the product of multiple separate transitions from sexual conspecifics, allowing us to treat distinct asexual lineages as replicated experiments into asexuality (Neiman *et al.* 2011; Paczesniak *et al.* 2013). The polyploidy (triploidy and tetraploidy; Wallace *et al.* 1992; Neiman *et al.* 2011) of asexual *P. antipodarum* also allows us to use comparisons between the two asexual ploidy levels and the diploid sexuals to study the effects of ploidy elevation.

What makes *P. antipodarum* particularly well suited for this study is that multiple asexual female lineages occasionally (~1.5% of all offspring, depending on the asexual lineage involved; Neiman *et al.* 2012) produce males, perhaps *via* loss of sex chromosomes during oogenesis (Soper *et al.* 2013). We leverage this unusual phenomenon to characterize morphological variation of sperm in sexually *vs*. asexually produced males, with the expectation that the latter have experienced a relatively recent history of relaxed selection on male-specific traits and genes. A general pattern of distinct phenotypes and, in particular, higher variation across sperm phenotypes from asexual *vs*. sexual males would be consistent with this prediction.

While we have been implicitly assuming that male genotype drives sperm phenotype (Hecht 1998), we also took advantage of the fact that eukaryotic cell size and nuclear genome content are typically positively correlated (Gregory 2001; Jovtchev *et al.* 2006; Beaulieu *et al.* 2008) - including in sperm cells (Cui 1997; Gallardo *et al.* 1999; van Munster *et al.* 1999)- to establish predictions that can also enable us to decouple effects of nuclear genome DNA content (which accounts for both ploidy elevation and diploid genome size changes) from those of transitions to asexuality. In particular, we expected that the mean size of sperm heads will increase with increasing nuclear genome content (regardless of reproductive mode), but that increased nuclear genome DNA content will not be associated with increased sperm tail length because the tail does not contain nuclear DNA. In other words, dual main effects of nuclear DNA content and reproductive mode on sperm morphology in *Potamopyrgus* should manifest as consistently larger sperm heads (nuclear DNA content) along with consistently higher variance in both sperm heads and tails in the asexually produced males.

## Materials and Methods

### Snail collection and ploidy/reproductive mode determination

*Potamopyrgus antipodarum* used in this study were collected either from New Zealand lake populations (7 lakes; 21 sexual males) or from individual and separately derived asexual lineages (8 lineages; 24 asexual males). The latter are recently descended (<5 generations) from asexual snails collected from New Zealand lake populations that were subsequently cultured in our laboratory at the University of Iowa. All snails were housed in plastic 10L tanks filled with oxygenated carbon-filtered tap water in a 16C room with a 16:8-hr light:dark cycle and fed dried *Spirulina* algae *ad libitum* (following Zachar & Neiman 2013). We focused our search for asexual males in lineages that had been previously characterized as relatively likely to harbor asexual males (Neiman *et al.* 2012) and asexual lineages descended from lake populations that population genetic studies (Paczesniak *et al.* 2013) demonstrated harbor asexual subpopulations that represent separate *in situ* transitions from sexual to asexual reproduction. For the sexual males, we used field collections from lakes that were known to contain relatively high frequencies (>10-20%) of sexual males (Neiman *et al.* 2011).

We identified males by turning each individual snail on its back (*i.e.*, foot up) under a dissecting microscope and then carefully looking for the penis, evident on the right side of the body of adult male snails while the snail is attempting to right itself. We used snails from field collections (sexed a total of 1370 snails) or lineages (sexed a total of 4505 snails) that were large enough (>/=3mm shell length) to have reached sexual maturity (*e.g.*, Larkin *et al.* 2016). Because we did not find any asexual males in field collections despite evaluating > 380 field-collected male *P. antipodarum* from eight lake populations (unsurprising in light of the scarcity of these males amongst all asexual *P. antipodarum*; Neiman *et al.* 2011, 2012), we had to compare field-collected sexual individuals to laboratory-reared asexual males. Because environmental conditions can influence sperm morphology (Perez-Crespo & Guitierrez-Adan 2008; Immler *et al.* 2010; Bakos *et al.* 2011), we addressed whether rearing conditions had an effect on sperm morphology by comparing 12 sperm from each of three males from a lab-reared sexual lineage to 12 sperm from each of three wild-caught sexual males from the same lake of origin (Alexandrina) as the lab-reared sexuals. There were no significant differences between lab-reared and field-collected males in any of the traits that we measured (Supp Figure 1A). A principal component analysis demonstrated nearly complete overlap between wild-caught and lab-reared snails when plotting the first two principal components (explaining 77.7% of all variation) of sperm traits (Supp Figure 1B), suggesting that field *vs*. lab rearing environments are not likely to provide a major source of variation in sperm morphology.

We used a Becton Dickinson ARIA II flow cytometer on dissected head tissue (following Bankers *et al.* 2017) to assign ploidy (diploid, triploid, tetraploid) and, thus, reproductive mode (diploid = sexual; triploid, tetraploid = asexual; Wallace 1992; Neiman *et al.* 2011) to each male. The overall incidence of males in asexual lineages was ~1% (59/5875), and asexual lineages varied in the relative frequency of male production (Supp Table 1). Both observations regarding male frequency in asexual lineages are consistent with a similar survey in Neiman *et al.* (2012). Because our experimental design goal was to replicate males three times within each asexual lineage that we used, and because we did not want to perform comparisons involving variation with unbalanced sample sizes, we haphazardly excluded 35 males that either represented a lineage in which more than three males were found (13 asexual lineages) or were from a lineage in which fewer than three males were found (six asexual lineages). This left us with a total of 24 asexually produced male *P. antipodarum* for our study. For sexual males, we used the first three haphazardly selected sexual males that we identified per each of seven field collections, for a total of 21 sexual male *P. antipodarum*.

Hereafter, we use “set” to indicate one three-male replicate from each unique lineage or lake of origin and ploidy combination: our diploid *P. antipodarum* sample represents seven male sets (each from a different field collection), triploid *P. antipodarum* are represented by six male sets (each from a different asexual lineage and lake of origin), and tetraploid *P. antipodarum* are represented by two male sets (from two different asexual lineages, each from a different lake), for a total of 15 *P. antipodarum* sets and 45 males (Figure 1).

**Figure 1.**
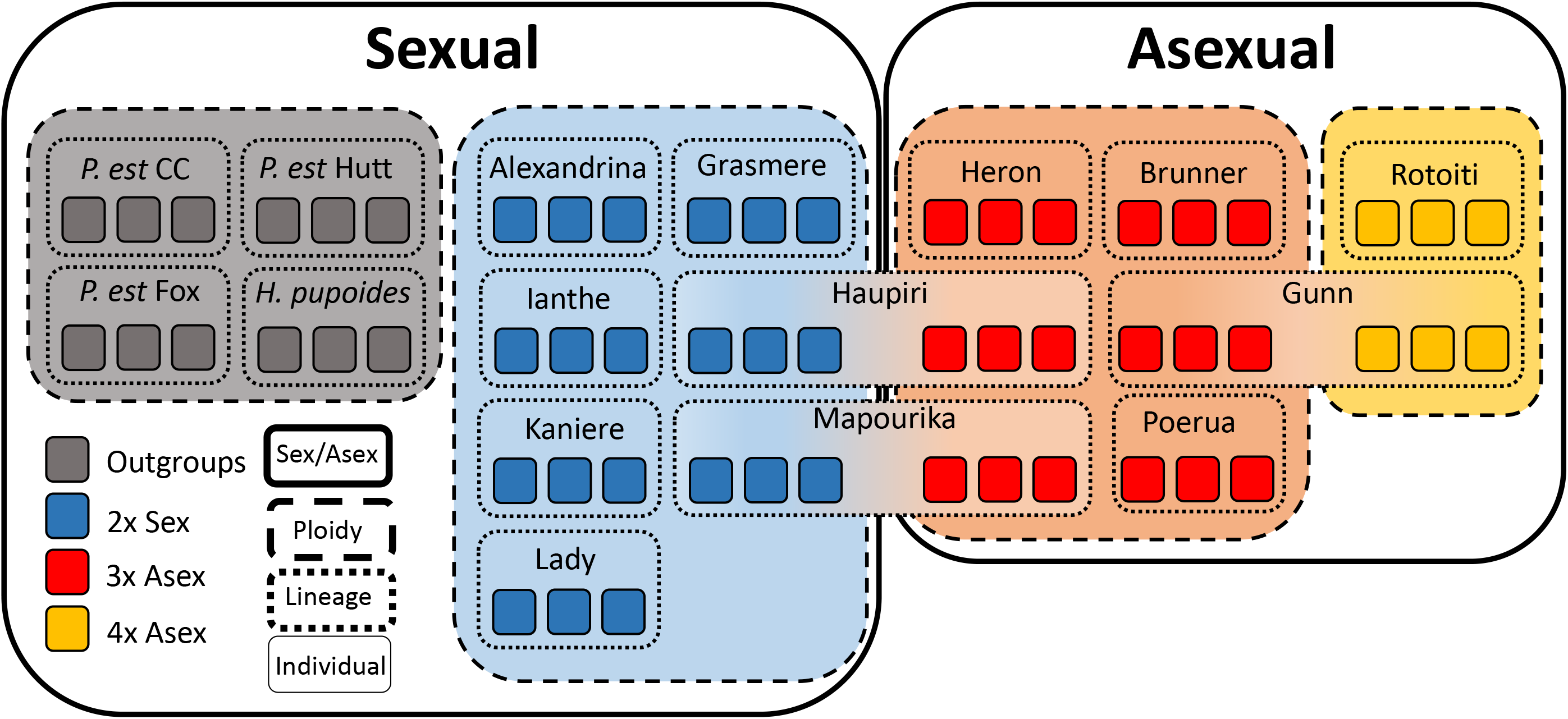
Schematic representing our experimental design, which included sperm from 57 individual males (each male represented by small colored boxes) from three different species, two reproductive modes, three ploidy levels, 12 lakes, and three estuaries. We characterized at least 12 sperm per male and three males per set, for a total of at least 36 sperm per set. Each set is represented by the boxed lake of origin and three small colored boxes representing the three males in the set. *P. est* = *P. estuarinus* from three different New Zealand estuaries (CC = Waikuku Beach; Hutt = Hutt River, NZ; Fox = Foxton). *H. pupoides* was collected from Waikuku Beach.

We provided a separate control for nuclear genome DNA content, which can have major influences on sperm phenotype (Alavi *et al.* 2013), by including males and sperm from the closely related (Haase 2008) and diploid obligate sexual species *P. estuarinus* (three sets, for a total of nine males) and *Halopyrgus pupoides* (one set, three males). Because *P. estuarinus* and *H. pupoides* have only about 60% of the nuclear genome DNA content of diploid *P. antipodarum* (Logsdon *et al.* 2017), we used comparisons across these three diploid taxa to account for the effect of nuclear genome DNA content (*vs*. ploidy and reproductive mode) on sperm phenotype. These outgroup snails were collected from their native New Zealand estuary habitat (Waikuku Beach, New Zealand) in January 2018.

### Sperm extraction and imaging

We used forceps to extract the bodies of male snails from their shells and then dissected away the sperm duct, which transports sperm and seminal fluid from the testis to the penis. We centered the sperm duct tissue on a glass microscope slide and added ~5μl of DAPI staining solution (following Soper *et al.* 2013). We used repeated pipetting of this solution to release sperm from the ruptured duct and distribute the sperm across the slide. We then mounted the slide on an Olympus IX71 fluorescence microscope and used the DAPI filter to locate the fluorescent sperm heads and the polarized filter to photograph sperm for subsequent analysis. We haphazardly selected and then photographed a minimum of 12 whole sperm cells for every male. We chose 12 sperm because the amount of variation around the mean when including an increasing amount of sperm measurements asymptotes at ~10 sperm in our rarefaction analysis (see Supp Figure 2). We excluded sperm cells that were entangled with other sperm or were obscured by non-sperm cellular detritus. We used sperm from at least three males (one male set) for each unique lake and ploidy combination, for a total of at least 36 sperm per set.

### Phenotype measurements

We imported photographs into ImageJ (Rueden *et al.* 2017) and measured head length (HL; head tip to head base), head width (HW; width at midpoint of head), head circumference (HC; perimeter of head), and tail length (TL; length from head base to tail end) for each sperm cell. We used our consistent observations of a number of qualitatively distinct phenotypic classes of sperm abnormalities to sort sperm into “normal” (most common phenotype of smooth cylindrical head and smooth tail of consistent width; 79% of all sperm) *vs*. one of six “abnormal” ultrastructural categories: >1 tail, distorted neck, oblong head, tail bump, two heads, wrinkled head (*e.g.*, Menkveld *et al.* 1990). This approach allows us to distinguish large-scale differences in complex sperm morphology that might not be evident from our measurements of individual sperm traits.

### Statistical analyses

We used a RAxML tree (CIPRES V. 3.1) generated from Illumina paired-end whole-genome sequencing data (70,437 SNPs) from 26 *P. antipodarum* (10 sexual individuals, 16 asexual individuals (11 triploid and five tetraploid)) to provide a phylogenetic framework needed to control for potential effects of phylogenetic relationship on sperm phenotype (Supp Figure 3). This tree also demonstrates that we have likely captured a broad sample of sexual and asexual *P. antipodarum* genetic diversity, including a number of separately derived asexual lineages. We used this tree in Phylosignal (Keck *et al.* 2016), an R package that tests for phylogenetic signal, to assess whether sperm morphology was determined at least in part by shared ancestry. We attached mean trait values for each of the sperm traits measured below to a subsampled phylogenetic tree, which represented directly (same asexual lineage) or indirectly (shared lake of origin, justified by multiple lines of evidence for a predominant effect of source population on *P. antipodarum* genotype, *e.g.*, Dybdahl & Lively 1995; Paczesniak *et al.* 2013; Bankers *et al.* 2017) nine of the 15 *P. antipodarum* sets from which we sampled males (*i.e.*, sets with both genetic data and sperm measurements). We did not find any significant phylogenetic correlations underlying these sperm traits (Supp Figure 4). Together, the outcome of these analyses indicate that phylogenetic relatedness is not a main driver of variation in sperm morphology. We also used hierarchical clustering of sperm morphology data to produce a cladogram that we used to assess whether phylogenetic relatedness is a main source of across-lineage/lake sperm variation (Figure 2, Supp Figure 5).

**Figure 2.**
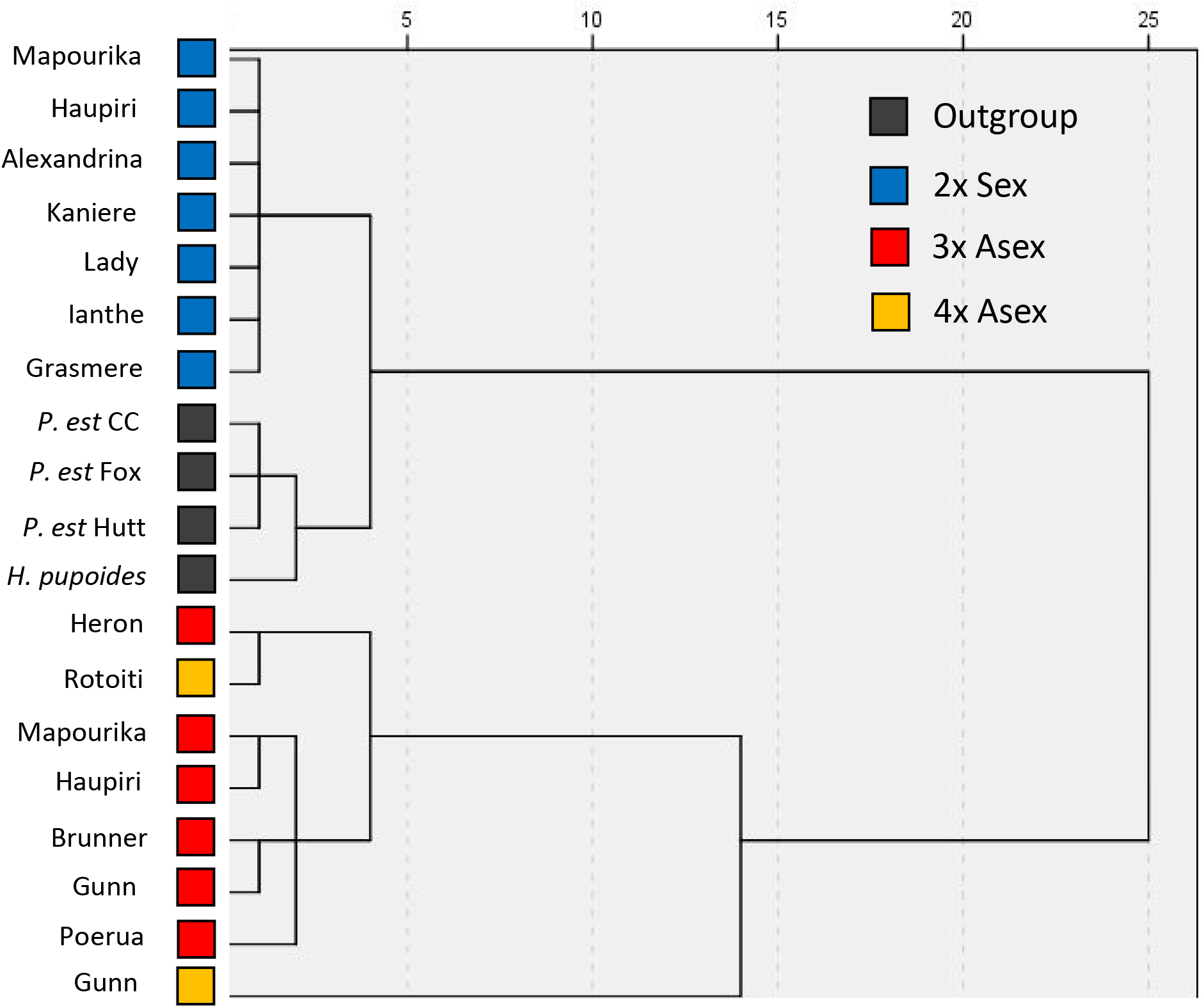
Hierarchical clustering dendrogram of sperm head morphology. Each square represents the mean trait values of the >/= 36 sperm taken from each of the three males used for each unique ploidy/reproductive mode/population set. Labeling follows Figure 1.

We then used the means calculated from the 12 sperm from each individual male for each trait (HL, HW, HC, TL) in the general linear model and Tukey posthoc pairwise comparison analysis framework as applied in SPSS 24 to assess whether the samples were analytically distinguishable across the comparison categories of nuclear genome DNA content, ploidy, and reproductive mode. Finally, we used Linear Discriminate Analysis (LDA) to assess whether and how ploidy level affects sperm morphology. This analysis approach minimizes within-group variance and maximizes across-group variance to find linear combinations of variables that best explain the data. All statistical analyses were implemented in R 3.4.1 (R Core Team 2013) or SPSS 24.0 (IBM Corp. 2016)

## Results

### Sperm morphology is primarily driven by nuclear genome content and reproductive mode

We evaluated a total of 750 sperm from each of 57 male snails (21 diploid sexuals, 18 triploid asexuals, six tetraploid asexuals, nine *P. estuarinus*, three *H. pupoides*) (Figure 1), derived from 12 lakes and three estuaries of origin as well as three species. Our analyses revealed extensive qualitative (Supp Figure 6) and quantitative variation (Supp Table 2, Supp Figure 7) in sperm morphology. Multi-dimensional hierarchical clustering of primary sperm traits (HL, HW, HC, TL) suggested that reproductive mode is a primary driver of sperm morphology in *P. antipodarum* (Figure 2). This conclusion is bolstered by the fact that diploid sexual *P. antipodarum* sperm clustered with the two diploid sexual outgroup species instead of with the asexual polyploid conspecifics.

We next used comparisons across the three ploidy levels represented by *P. antipodarum* and sperm from the diploid outgroup species *P. estuarinus* and *H. pupoides* to decouple effects of nuclear genome DNA content from ploidy and reproductive mode on sperm morphology. These comparisons revealed that nuclear genome DNA content seems to have a major effect on head shape, with head shape metrics increasing consistently with nuclear genome content. Nevertheless, that nuclear genome DNA content *per se* is not the sole driver of sperm morphology in these snails is demonstrated by several lines of evidence. First, sperm produced by *H. pupoides* and *P. estuarinus*, which seem to share the same relatively small nuclear DNA content (Logsdon *et al.* 2017), were nevertheless morphologically distinct (Figure 2, Supp Figure 9). Second, head width and tail length were statistically indistinguishable between triploid and tetraploid *P. antipodarum* (Supp Figure 7). Third, the CVs for triploid and tetraploid *P. antipodarum* are statistically indistinguishable for all head-related traits (Supp Figure 8). Fourth, while tail length is similar across different nuclear DNA contents in *P. antipodarum*, the coefficient of variation of our tail length estimates is significantly increased in triploid (CV mean = 12.2) and tetraploid (CV mean = 14.9) sets relative to diploids (CV mean = 4.7) (Supp Figure 8). Finally, our linear discriminant analyses demonstrate that the three different *P. antipodarum* ploidies partially overlap each other in morphometric space (in particular, triploid-produced sperm share space with both diploid and tetraploid-produced sperm) (Figure 3). We also detected relatively minor evidence for ploidy effects on sperm trait means and variances: the LDA revealed ploidy-based clustering in *P. antipodarum* (Figure 3), and the relative magnitude of change between measurements of *P. antipodarum* head length and head circumference across ploidy levels show a pattern consistent with a positive effect of ploidy-driven variation of nuclear genome DNA content on sperm head size.

**Figure 3.**
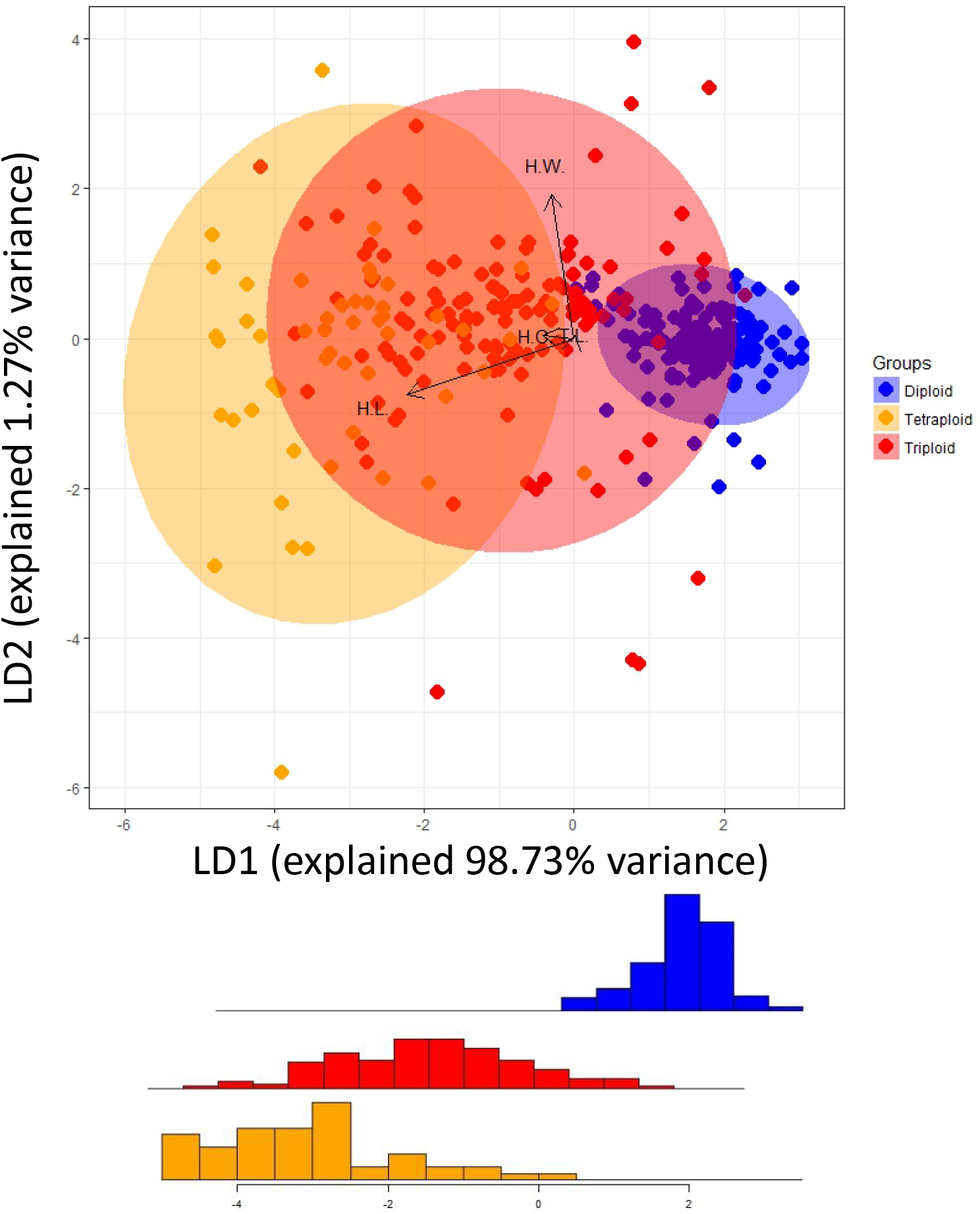
Linear discriminant function analysis of transformed sperm traits (head length, head width, head circumference, tail length) for diploid, triploid, and tetraploid *P. antipodarum*. Histogram represents the cumulative numbers of sperm in each bin for LD1. Arrows represent each variable (HL, HW, HC, TL), length of arrows represents degree of discrimination, and arrow angle is the direction of the variables in 2-D space. Shaded area encompasses 95% of data.

With respect to the latter point, the magnitude of fold difference in sperm head size traits is the lowest in the triploid/tetraploid comparison (*i.e.*, a 33% increase in nuclear genome DNA content), and the highest in the diploid/tetraploid comparison (*i.e.*, a 100% increase in nuclear genome DNA content), with the diploid/triploid values in between (*i.e.*, a 50% increase in nuclear genome DNA content). This pattern is as expected in a situation where sperm heads increase in size as nuclear DNA content increases, but only in proportion to the proportional increase in nuclear DNA content (Supp Figure 10).

Finally, as predicted under a scenario where the absence of sex translates into relaxed selection, CV-based analyses indicated that sperm produced by diploid male *P. antipodarum* harbors significantly lower variance (*e.g.*, HL mean = 4.48μm, SD = 0.09) than sperm produced by triploid (HL mean = 5.29μm, SD = 0.32) and tetraploid (HL mean = 5.99μm, SD = 0.22) males (Supp Figure 8).

### Asexual *P. antipodarum* often produce qualitatively abnormal sperm

Visual inspection of *P. antipodarum* sperm revealed substantial variation. Most (N=429; 76%) of these sperm had an overtly normal phenotype, represented by a smooth and cylindrical head and a tail of consistent width from the head base to tip (Supp Figure 6A). We sorted the most common deviations we observed from this normal phenotype into six discrete categories that emerged during our imaging and analysis stages: >1 tail (3.5% of all sperm), distorted neck (2.5%), oblong head (1.5%), tail bump (10%), two heads (1%), and wrinkled head (6.5%) (see Supp Figure 6B-D). There was markedly higher incidence of abnormal sperm produced by triploid (N=95; 43% of all triploid sperm) and tetraploid (N=24; 27%) *P. antipodarum* relative to the sperm produced by diploid male *P. antipodarum* (N=16; 6%) (Fisher’s exact test *p* = < 0.001) (Figure 4). Misshapen heads were the only abnormality shared across all three ploidies, while the most common abnormality, tail bump, is restricted to triploid asexual *P. antipodarum*.

**Figure 4.**
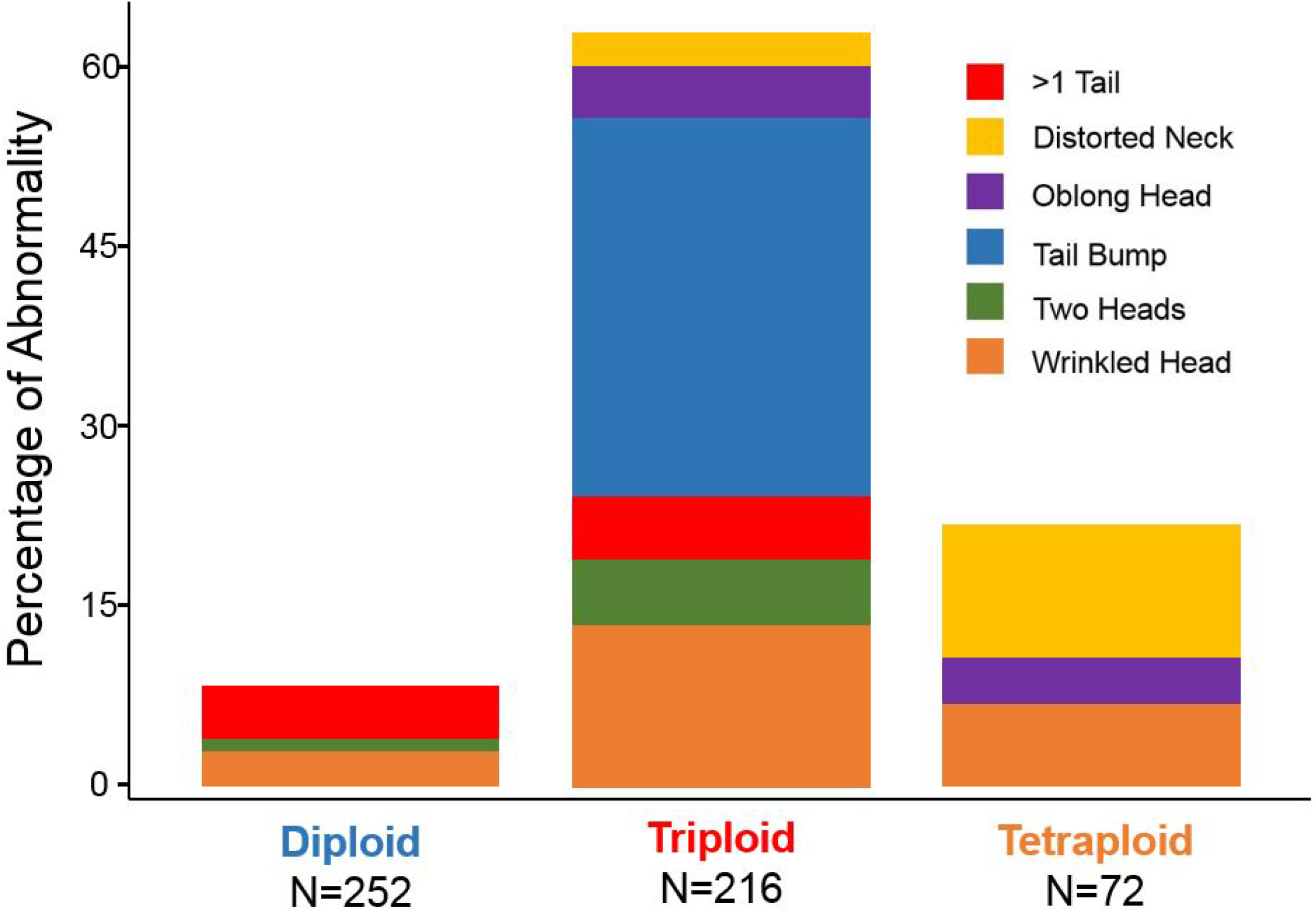
Proportion and characterization of sperm phenotypes that we categorized as “abnormal”.

### Among- and within-species variation in sperm morphology

We used a principal component analysis to visualize and compare multivariate sperm phenotypes across sexual *P. antipodarum* and the two closely related sexual outgroup species. This analysis revealed a distinct species-level signal, with no overlap between the clusters of sperm data corresponding to each species (Supp Figure 9). Our general linear model and Tukey posthoc analyses also demonstrated statistically significant differences in each of the four individual sperm traits amongst these three species (Supp Table 2).

## Discussion

We used the powerful *Potamopyrgus antipodarum* system to perform a direct comparison of male-specific traits across otherwise similar male snails that differ in ploidy level and reproductive mode. Our study is especially novel in comparing sperm phenotypes across reproductive modes as well as ploidy level and diploid nuclear genome sizes, allowing us to at least in part decouple effects of polyploidy and genome size from those of asexuality.

### Nuclear DNA content affects sperm head size, while reproductive mode affects sperm variation

We used comparisons across the three different *P. antipodarum* ploidies and the three different diploid sexual taxa included in our study to assess the extent to which increased nuclear genome DNA content in the asexual *P. antipodarum*, *versus* asexuality *per se*, might affect sperm morphology. That asexual-produced sperm are generally larger, and that these size differences to some extent scale with ploidy and nuclear genome DNA content (Supp Figure 7), is consistent with the former possibility. Even so, the significant differences in sperm trait values (*e.g.*, head length) between *P. estuarinus* and *H. pupoides*, which have virtually identical genome sizes (Logsdon *et al.* 2017), and the significantly increased tail length variation in asexuals indicates that other factors besides nuclear genome content (including but not necessarily limited to reproductive mode) are also important in determining sperm morphology. The different outcomes between the analyses of head *vs*. and tail-related traits also provide a strong line of evidence in support of our conclusion that both nuclear genome DNA content and reproductive mode contribute to *P. antipodarum* sperm morphology. In particular, we found that sperm head trait values (*e.g.*, mean HL diploid = 4.48 μm, SD = 0.09; triploid = 5.29 μm, SD = 0.32; tetraploid = 5.99 μm, SD =0.22) but not tail length (mean tail length diploid = 128.6 μm, SD = 3.34; triploid = 137.73, μm SD = 22.05; tetraploid = 137.52 μm, SD = 12.99) increase with genome size. This result is as predicted if nuclear genome DNA content affects the sperm nucleus size, which is housed in the sperm head. The fact that the CVs are distinctly higher for both head and tail traits in the asexual *vs*. sexual males (*e.g.*, CV mean HL diploid = 3.2, SD = 0.92; triploid = 5.8 SD, = 1.98; tetraploid = 5.5, SD = 1.66; CV mean TL diploid = 4.7, SD = 3.52; triploid = 12.2, SD = 10.41; tetraploid = 14.9, SD = 7.90), along with the pronounced increase of incidence of tail abnormalities in asexuals relative to sexuals, also points to a distinct role for reproductive mode. In particular, the higher CVs across both heads and tails in asexual males are as expected if relaxed selection on sperm traits is driving a higher rate of morphological evolution.

We can draw parallels and contrasts with the few other ploidy-focused sperm studies of which we are aware: Alavi *et al.* (2013) showed that tetraploid European weatherfish (*Misgurnus fossilis*) produce sperm that are 4.8% larger than sperm produced by their triploid counterparts, and Flajshans *et al.* (2008) found that triploid male Prussian carp (*Carassius gibelio*) make sperm with higher volume than sperm generated by diploid and tetraploid males. Considered together along with these two earlier studies, we can conclude that nuclear genome DNA content is not the sole driver of the patterns we observed in *P. antipodarum*.

### *Asexual* P. antipodarum *are characterized by abnormal sperm*

We found evidence for a major effect of sexual *vs*. asexual reproduction on sperm morphology, with sperm produced by asexual males harboring substantially more variation and much more likely to feature large-scale structural abnormalities (Figure 3, 4). That a substantial fraction of asexual-produced sperm is abnormal is consistent with a scenario where male-specific traits are no longer useful in an asexual lineage. The abnormal phenotypes that often characterized sperm from asexual male *P. antipodarum* do appear to be affected in part by ploidy: sperm from triploid asexual males spanned a wider range of abnormal phenotypes and were more than twice as likely to be abnormal than sperm from tetraploid counterparts. Cytogenetic characterization of some components of spermatogenesis in diploid sexual *vs*. triploid asexual *P. antipodarum* have demonstrated that sperm production in the latter is associated with aberrant meiotic pairing (Soper *et al.* 2013). Meiosis is disrupted when odd number of chromosomes pair (Gupta 1985; Comai 2005; Henry *et al.* 2005), hinting that meiotic disruption and the across-sperm variation in genome size and gene content that this disruption might cause is one possible explanation for the differences between triploid and tetraploid-produced sperm in *P. antipodarum*.

There are at least two other non-mutually exclusive potential explanations for the increased frequency of production of abnormal sperm in asexual *vs*. snails. One possibility is mutation accumulation in the genes underlying sperm production, which are expected to be under relaxed selection in all (or nearly all)-female asexual populations (Schurko & Logsdon 2008; Lahti *et al.* 2009). Under this scenario, genes that influence sperm traits in asexuals should be accumulating mutations at a higher rate than sexuals. This mutation accumulation process would be evidenced by the degeneration and eventual pseudogenization of genes and traits related to sperm function in asexual populations. Second, it is possible that the increased incidence of sperm abnormalities in asexuals is affected by the higher incidence of aneuploidy in the asexually produced males (Neiman *et al.* 2011, Soper *et al.* 2013). While we cannot formally exclude this possibility, we would expect that aneuploidy-linked deformities would also be evident in other aspects of asexual male phenotype. Counter to this expectation, asexual male *P. antipodarum* are to the extent that we know behaviorally and morphologically indistinguishable from sexual males (*e.g.*, Soper *et al.* 2015) except in their sperm traits.

The fact that the male offspring occasionally produced by female asexual *P. antipodarum* are still investing in sperm suggests that the time period over which the genes underlying sperm production have been subject to relaxed selection is relatively short and/or that selection favoring the loss of this trait is weak. The latter would not be surprising in light of the fact that male production is quite rare (~<1.5% of all offspring) in asexual *P. antipodarum*. A powerful test of the former could come from comparisons of males produced by different asexual *P. antipodarum* lineages, which are often recently derived (Paczesniak *et al.* 2013) but also vary in the time since derivation from sexual *P. antipodarum* (Neiman *et al.* 2005). If time since derivation to asexuality is a key determinant of the extent of male trait degeneration, we predict that males from older asexual lineages should tend to produce sperm that harbors relatively high variance and is abnormal at a higher frequency than sperm produced by males from younger asexual lineages.

While we cannot formally exclude the possibility that ploidy elevation *per se* is a major driver of sperm abnormality, the fact that diploid male sexual *P. antipodarum* produce normal sperm at a similar frequency to the two outgroup species suggests that nuclear genome DNA content alone does not account for the high frequency of sperm abnormalities observed in asexual male *P. antipodarum*. The wide sperm phenotype variation that we observed across asexual males also hints at a process (*i.e.* mutation accumulation) that is stochastic compared to a direct consequence of ploidy elevation.

## Summary & Conclusion

Together, our results indicate that asexual male *P. antipodarum* are likely experiencing decay of male-specific traits, highlighting the likelihood that transitions to asexual reproduction results in striking differences in selective regimes between sexual and asexual populations. This finding is of direct relevance to evaluating how relaxed selection influences phenotypes in natural populations and suggests that transitions back to sexual reproduction in asexual populations are likely to face challenges associated with degeneration of sex-specific traits. Critical next steps include the characterization of molecular evolution of sperm-related genes in sexual *vs*. asexual populations and assessment of whether older asexual populations have experienced a relatively high level of trait decay.

## Acknowledgments

We gratefully acknowledge help in snail collections from Mike Winterbourn, Mary Morgan-Richards, Katelyn Larkin, Kyle McElroy, Jeremy Richardson, Laura Bankers, and Kaitlin Hatcher. We thank Sarit Smolikove for microscope use, Julie Meachen for statistical advice, and Deanna Soper for assistance in sperm extraction protocols. Three anonymous reviewers provided very helpful feedback. The ploidy identification data presented herein were obtained at the Flow Cytometry Facility, which is a Carver College of Medicine/Holden Comprehensive Cancer Center core research facility at the University of Iowa. The Facility is funded through user fees and the generous financial support of the Carver College of Medicine, Holden Comprehensive Cancer Center, and Iowa City Veteran’s Administration Medical Center. This project was funded by NSF-MCB 1122176 and NSF DEB-1753851

