## Supplemental Material for "Male evolution under relaxed selection: Evidence for degeneration in sperm produced by male snails from asexual lineages"

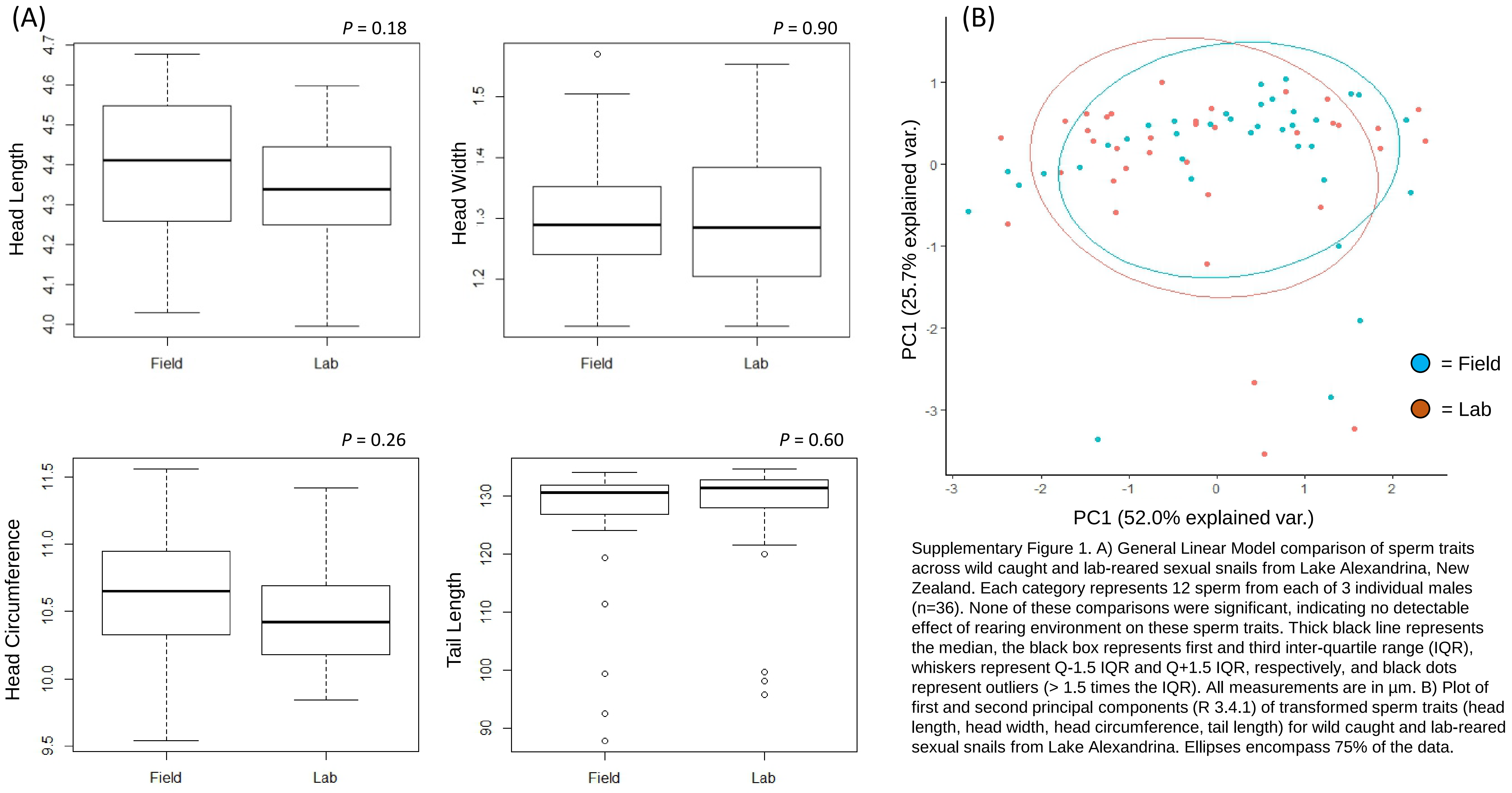


Supplementary Figure 1. A) General Linear Model comparison of sperm traits across wild caught and lab-reared sexual snails from Lake Alexandrina, New Zealand. Each category represents 12 sperm from each of 3 individual males (n=36). None of these comparisons were significant, indicating no detectable effect of rearing environment on these sperm traits. Thick black line represents the median, the black box represents first and third inter-quartile range (IQR), whiskers represent Q-1.5 IQR and Q+1.5 IQR, respectively, and black dots represent outliers (> 1.5 times the IQR). All measurements are in µm. B) Plot of first and second principal components (R 3.4.1) of transformed sperm traits (head length, head width, head circumference, tail length) for wild caught and lab-reared sexual snails from Lake Alexandrina. Ellipses encompass 75% of the data.

| Lake | Source | 2x | 3x | 4x | N | % | Lat | Long |
|---|---|---|---|---|---|---|---|---|
| Alexandrina | Field | 3 | 0 | 0 | n/a | n/a | -43.9401 | 170.4484 |
| Brunner | Lab | 0 | 3 | 0 | 250 | 1.2 | -42.6169 | 171.4416 |
| Grasmere | Field | 3 | 0 | 0 | 200 | 0.0 | -41.7311 | 174.1685 |
| Gunn | Lab | 0 | 14 | 10 | 840 | 2.8 | -44.8744 | 168.0911 |
| Haupiri | Field/Lab | 3 | 4 | 0 | 320 | 1.2 | -42.5671 | 171.6889 |
| Heron | Lab | 0 | 9 | 0 | 575 | 1.5 | -43.4739 | 171.1792 |
| Ianthe | Field | 3 | 0 | 0 | n/a | n/a | -43.0533 | 170.6194 |
| Kaniere | Field | 3 | 0 | 0 | 80 | 0.0 | -42.8428 | 171.1443 |
| Lady | Field | 3 | 0 | 0 | n/a | n/a | 42.6030 | 171.5739 |
| Mapourika | Field/Lab | 3 | 4 | 0 | 520 | 0.7 | -43.3165 | 170.1973 |
| Moeraki | Lab | 0 | 1 | 0 | 230 | 0.4 | -43.7297 | 169.2805 |
| Okareka | Lab | 0 | 0 | 0 | 275 | 0.0 | -38.1720 | 176.3624 |
| Poerua | Lab | 0 | 6 | 4 | 1045 | 0.9 | -42.7054 | 171.4931 |
| Rotoiti | Lab | 0 | 0 | 4 | 310 | 1.2 | -38.0317 | 176.4196 |
| Selfe | Field | 0 | 0 | 0 | 100 | 0.0 | -43.2387 | 171.5174 |
| Taupo | Lab | 0 | 0 | 0 | 130 | 0.0 | -38.7923 | 175.9057 |
| Taylor | Lab | 0 | 0 | 0 | 200 | 0.0 | -42.7667 | 172.2302 |
| Te Anau | Lab | 0 | 0 | 0 | 200 | 0.0 | -45.2703 | 167.7517 |
| Waikaremoana | Lab | 0 | 0 | 0 | 600 | 0.0 | -38.7657 | 177.0935 |
| Total | | 21 | 41 | 18 | 5875 | 1.3 | | |

Supplementary Table 1. Summary of sampling scheme. “Lake” = original source of the male (field collections), or the ancestors of the male (laboratory lineages) from New Zealand. “2x”, “3x”, “4x” represents the total number of males of each ploidy type found in the collection or lineage as whole, respectively. N refers to the total number of snails sexed for each collection or lineage; % = percentage of that collection/lineage that was male. Source = field vs. laboratory lineages for a particular lake. “Lat” = latitude and “Long” = longitude for each lake. “n/a” represents populations or lineages that harbor >~10% males and thus harbor a relatively high frequency of sexual individuals.

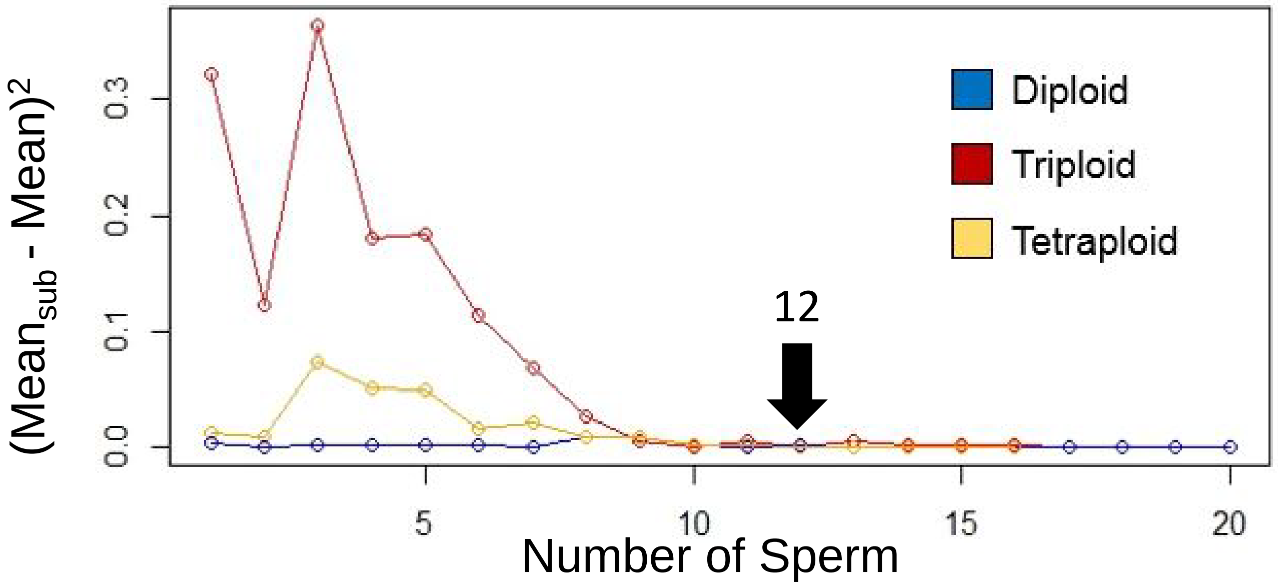


Supplementary Figure 2. Rarefaction analysis to assess sampling number on capturing variation of sperm size. We used randomly selected sperm from diploid, triploid and, and tetraploid males to assess how the number of sperm measured reflects lineage variation. We first determined the overall mean from each category. We then took the mean of two randomly selected sperm and subtracted it from the overall mean and squared it. A value of ~zero indicates that the overall mean and mean of the subsampled sperm are indistinguishable. We repeated this process to 20 sperm and found that 12 sperm sufficiently captures the variation from each category (*i.e.*, at the point where the number of sperm included in the analysis generates an asymptote at 0).

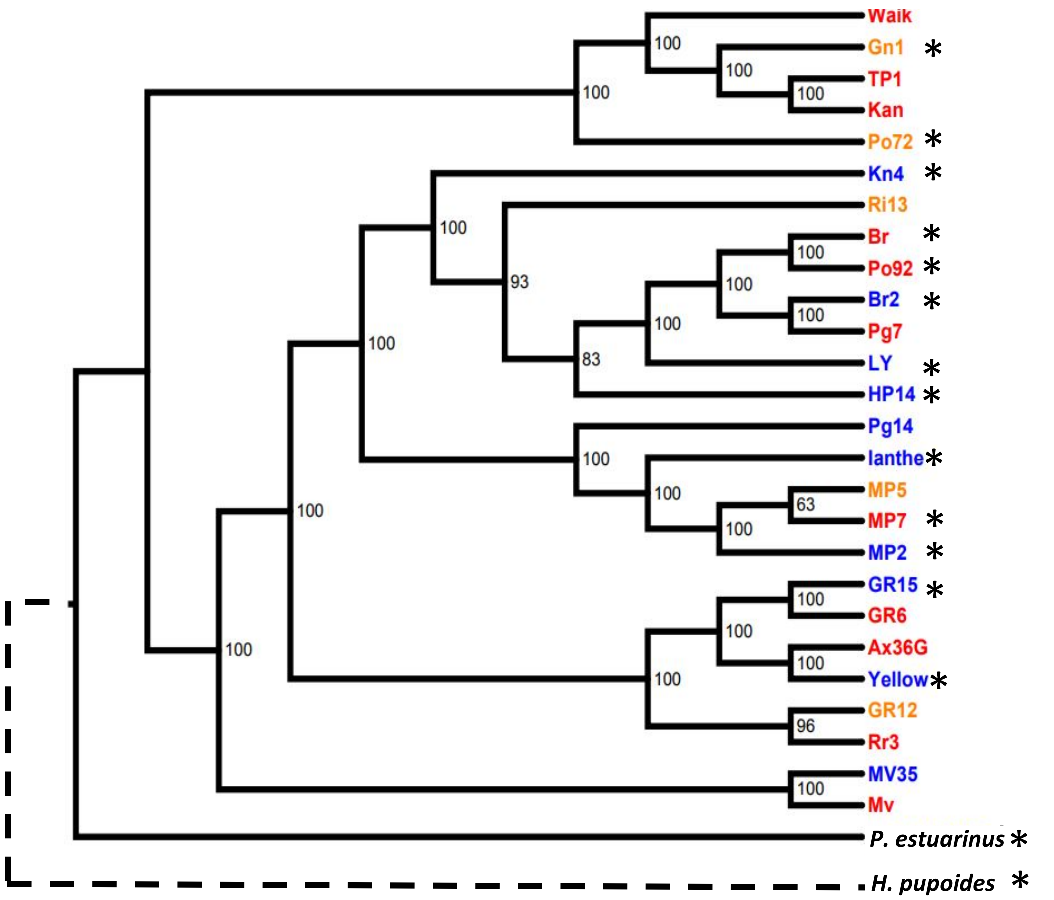


Supplementary Figure 3. RAxML genetic tree built from Illumina whole-genome SNP data (available upon request). Blue = diploid, red = triploid, orange = tetraploid. Asterisk indicates linages from which sperm was sampled. Dotted line reflects the predicted relationship of *H. pupoides* to these samples based on the mitochondrial tree presented in Haase (2008). Numbers indicate bootstrap support for each node based on 500 comparisons.

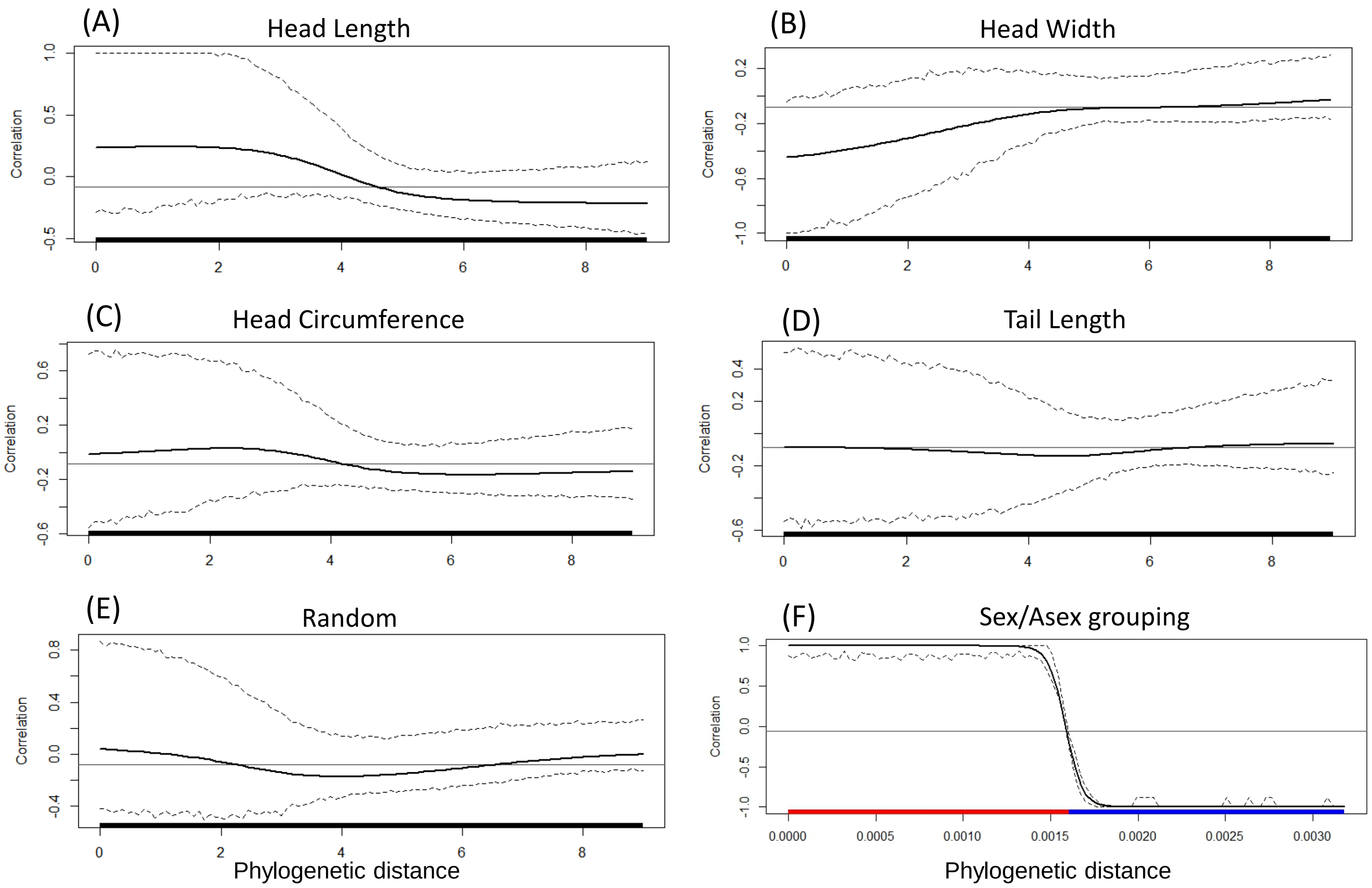


Supplementary Figure 4. Phylogenetic correlogram (Keck *et al.* 2016) for sperm traits (A-D), random sampling (E), and forcing the phylogenetic grouping of sexual and asexual individuals (F). The horizontal black line is the expected Moran's I given the null hypothesis of no phylogenetic autocorrelation. Solid black line indicates Moran's I index of autocorrelation, and dashed lines indicate the bounds of the 95% confidence interval of Moran's I. The thick bar at the bottom of each plot represents statistical significance, or lack thereof, of phylogenetic autocorrection: red indicates significantly positive autocorrelation, blue indicates significantly negative autocorrelation, and black represents no significant autocorrelation. Trait data (A-D) show no statistical differences from random phylogenetic placement (E).

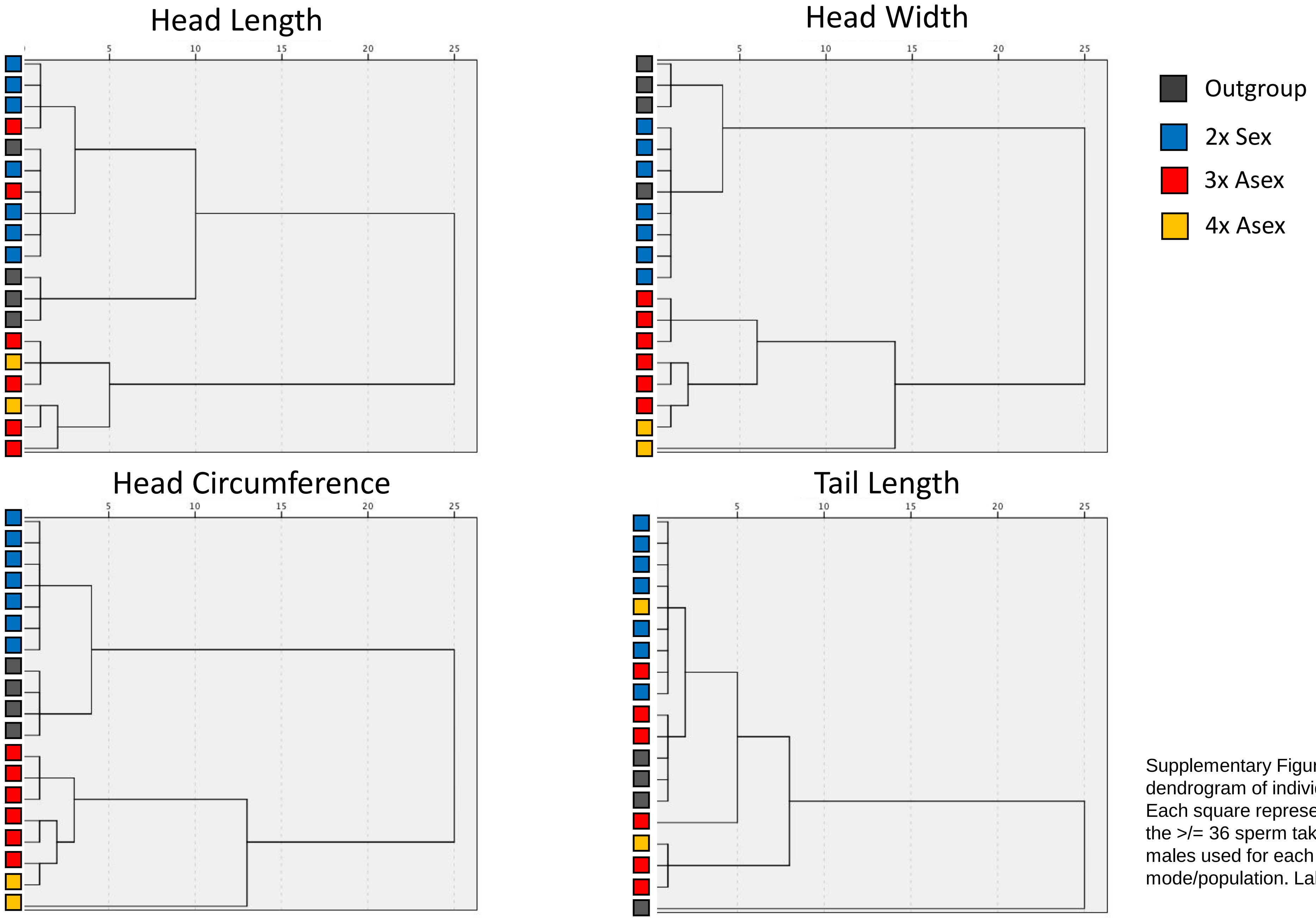


Supplementary Figure 5. Hierarchical clustering dendrogram of individual sperm trait morphology. Each square represents the mean trait values of the >/= 36 sperm taken from each of the three males used for each unique ploidy/reproductive mode/population. Labeling follows Fig. 1.

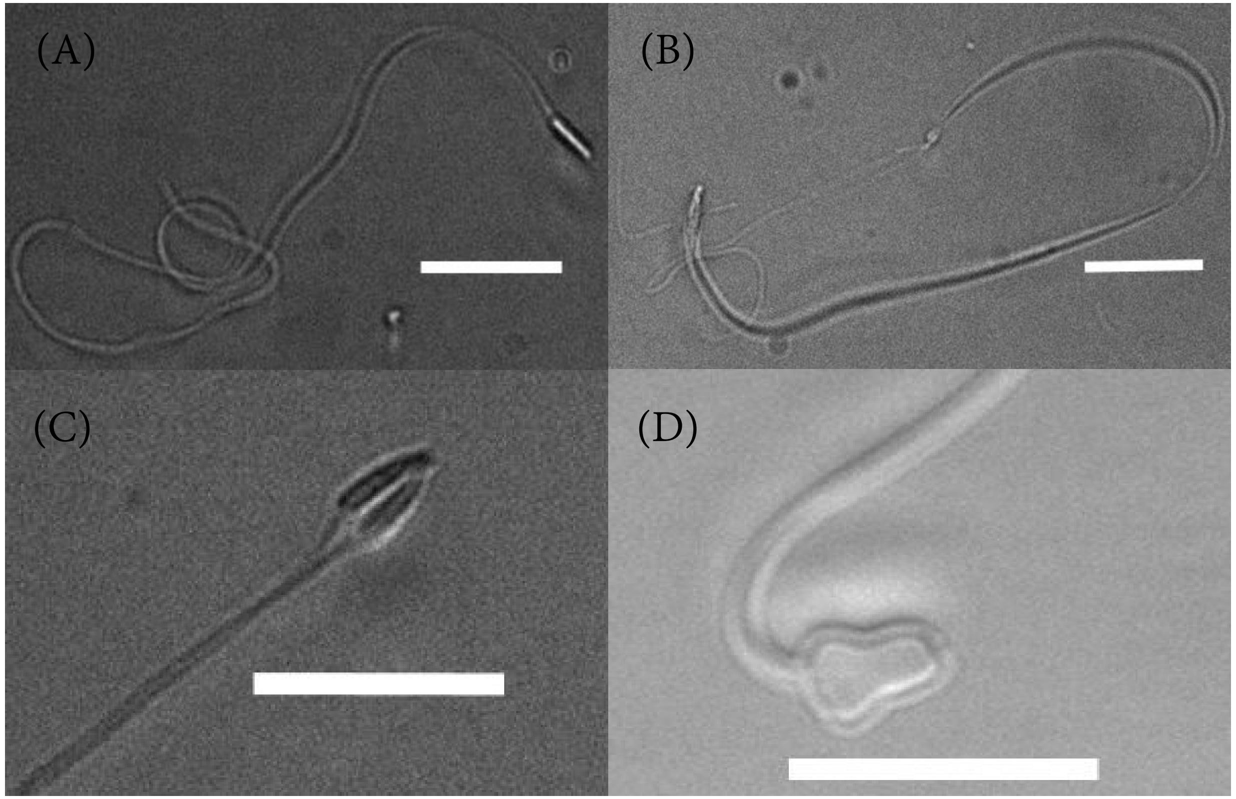


Supplementary Figure 6. Sperm ultrastructure variation in *P. antipodarum*. Images not to scale. (A) Common “normal” phenotype, (B) Two-tailed, (C) Two-headed, (D) Oblong. White bar scaled to 10.71µm.

## HL

| (I) set | (J) set | Mean Difference (I-J) | Std. Error | Sig. | 95% Confidence Interval | |
|---|---|---|---|---|---|---|
| | | | | | Lower Bound | Upper Bound |
| 1-H.pup | 2-P.est | .2663131313 | .1381375091 | .316 | -.124034670 | .6566609328 |
| | 3-Diploid | -.156952551 | .1278904824 | .736 | -.518344391 | .2044392898 |
| | 4-Triploid | -.961890912* | .1292158078 | .000 | -1.32702785 | -.596753978 |
| | 5-Tetraploid | -1.66050097* | .1465169541 | .000 | -2.07452734 | -1.24647461 |
| 2-P.est | 1-H.pup | -.266313131 | .1381375091 | .316 | -.656660933 | .1240346702 |
| | 3-Diploid | -.423265682* | .0825529514 | .000 | -.656543112 | -.189988252 |
| | 4-Triploid | -1.22820404* | .0845916029 | .000 | -1.46724228 | -.989165809 |
| | 5-Tetraploid | -1.92681410* | .1092072897 | .000 | -2.23541114 | -1.61821707 |
| 3-Diploid | 1-H.pup | .1569525507 | .1278904824 | .736 | -.204439290 | .5183443913 |
| | 2-P.est | .423265682* | .0825529514 | .000 | .1899882520 | .6565431120 |
| | 4-Triploid | -.804938361* | .0665563172 | .000 | -.993012638 | -.616864085 |
| | 5-Tetraploid | -1.50354842* | .0959178618 | .000 | -1.77459230 | -1.23250454 |
| 4-Triploid | 1-H.pup | .961890912* | .1292158078 | .000 | .5967539783 | 1.327027846 |
| | 2-P.est | 1.22820404* | .0845916029 | .000 | .9891658094 | 1.467242277 |
| | 3-Diploid | .804938361* | .0665563172 | .000 | .6168640846 | .9930126381 |
| | 5-Tetraploid | -.698610059* | .0976779694 | .000 | -.974627637 | -.422592482 |
| 5-Tetraploid | 1-H.pup | 1.66050097* | .1465169541 | .000 | 1.246474605 | 2.074527337 |
| | 2-P.est | 1.92681410* | .1092072897 | .000 | 1.618217069 | 2.235411136 |
| | 3-Diploid | 1.50354842* | .0959178618 | .000 | 1.232504540 | 1.774592301 |
| | 4-Triploid | .698610059* | .0976779694 | .000 | .4225924817 | .9746276367 |

Based on observed means.
The error term is Mean Square(Error) = .043.
*. The mean difference is significant at the 0.05 level.

## HW

| (I) set | (J) set | Mean Difference (I-J) | Std. Error | Sig. | 95% Confidence Interval | |
|---|---|---|---|---|---|---|
| | | | | | Lower Bound | Upper Bound |
| 1-H.pup | 2-P.est | .0013005051 | .1417823054 | 1.000 | -.399346731 | .4019477412 |
| | 3-Diploid | -.186438970 | .1312649082 | .618 | -.557366234 | .1844882947 |
| | 4-Triploid | -.409636746* | .1326252026 | .026 | -.784407919 | -.034865574 |
| | 5-Tetraploid | -.500193085* | .1503828444 | .013 | -.925143651 | -.075242518 |
| 2-P.est | 1-H.pup | -.001300505 | .1417823054 | 1.000 | -.401947741 | .3993467311 |
| | 3-Diploid | -.187739475 | .0847311339 | .190 | -.427171994 | .0516930449 |
| | 4-Triploid | -.410937252* | .0868235757 | .000 | -.656282575 | -.165591928 |
| | 5-Tetraploid | -.501493590* | .1120887543 | .000 | -.818233041 | -.184754139 |
| 3-Diploid | 1-H.pup | .1864389695 | .1312649082 | .618 | -.184488295 | .5573662337 |
| | 2-P.est | .1877394746 | .0847311339 | .190 | -.051693045 | .4271719941 |
| | 4-Triploid | -.223197777* | .0683124241 | .016 | -.416234445 | -.030161108 |
| | 5-Tetraploid | -.313754115* | .0984486811 | .020 | -.591949563 | -.035558667 |
| 4-Triploid | 1-H.pup | .409636746* | .1326252026 | .026 | .0348655738 | .7844079192 |
| | 2-P.est | .410937252* | .0868235757 | .000 | .1655919276 | .6562825754 |
| | 3-Diploid | .223197777* | .0683124241 | .016 | .0301611084 | .4162344455 |
| | 5-Tetraploid | -.090556338 | .1002552296 | .894 | -.373856716 | .1927440393 |
| 5-Tetraploid | 1-H.pup | .500193085* | .1503828444 | .013 | .0752425183 | .9251436511 |
| | 2-P.est | .501493590* | .1120887543 | .000 | .1847541386 | .8182330409 |
| | 3-Diploid | .313754115* | .0984486811 | .020 | .0355586670 | .5919495633 |
| | 4-Triploid | .0905563382 | .1002552296 | .894 | -.192744039 | .3738567158 |

Based on observed means.
The error term is Mean Square(Error) = .045.
*. The mean difference is significant at the 0.05 level.

## HC

| (I) set | (J) set | Mean Difference (I-J) | Std. Error | Sig. | 95% Confidence Interval | |
|---|---|---|---|---|---|---|
| | | | | | Lower Bound | Upper Bound |
| 1-H.pup | 2-P.est | .5129326599 | .2965380793 | .425 | -.325022130 | 1.350887449 |
| | 3-Diploid | -.653739453 | .2745409142 | .137 | -1.42953484 | .1220559334 |
| | 4-Triploid | -2.53060501* | .2773859737 | .000 | -3.31443994 | -1.74677008 |
| | 5-Tetraploid | -3.98592881* | .3145261301 | .000 | -4.87471408 | -3.09714354 |
| 2-P.est | 1-H.pup | -.512932660 | .2965380793 | .425 | -1.35088745 | .3250221295 |
| | 3-Diploid | -1.16667211* | .1772153981 | .000 | -1.66744588 | -.665898344 |
| | 4-Triploid | -3.04353767* | .1815917459 | .000 | -3.55667809 | -2.53039726 |
| | 5-Tetraploid | -4.49886147* | .2344339359 | .000 | -5.16132289 | -3.83640004 |
| 3-Diploid | 1-H.pup | .6537394534 | .2745409142 | .137 | -.122055933 | 1.429534840 |
| | 2-P.est | 1.16667211* | .1772153981 | .000 | .6658983444 | 1.667445882 |
| | 4-Triploid | -1.87686556* | .1428756216 | .000 | -2.28060228 | -1.47312884 |
| | 5-Tetraploid | -3.33218935* | .2059056856 | .000 | -3.91403589 | -2.75034281 |
| 4-Triploid | 1-H.pup | 2.53060501* | .2773859737 | .000 | 1.746770080 | 3.314439944 |
| | 2-P.est | 3.04353767* | .1815917459 | .000 | 2.530397256 | 3.556678087 |
| | 3-Diploid | 1.87686556* | .1428756216 | .000 | 1.473128838 | 2.280602278 |
| | 5-Tetraploid | -1.45532380* | .2096840868 | .000 | -2.04784731 | -.862800282 |
| 5-Tetraploid | 1-H.pup | 3.98592881* | .3145261301 | .000 | 3.097143536 | 4.874714078 |
| | 2-P.est | 4.49886147* | .2344339359 | .000 | 3.836400039 | 5.161322895 |
| | 3-Diploid | 3.33218935* | .2059056856 | .000 | 2.750342814 | 3.914035894 |
| | 4-Triploid | 1.45532380* | .2096840868 | .000 | .8628002818 | 2.047847310 |

Based on observed means.
The error term is Mean Square(Error) = .198.
*. The mean difference is significant at the 0.05 level.

## TL

| (I) set | (J) set | Mean Difference (I-J) | Std. Error | Sig. | 95% Confidence Interval | |
|---|---|---|---|---|---|---|
| | | | | | Lower Bound | Upper Bound |
| 1-H.pup | 2-P.est | -35.4847062* | 9.078207933 | .002 | -61.1378292 | -9.83158325 |
| | 3-Diploid | -29.3705284* | 8.404787374 | .008 | -53.1207053 | -5.62035149 |
| | 4-Triploid | -38.4966776* | 8.491885943 | .000 | -62.4929769 | -14.5003783 |
| | 5-Tetraploid | -38.2850794* | 9.628893586 | .002 | -65.4943253 | -11.0758336 |
| 2-P.est | 1-H.pup | 35.4847062* | 9.078207933 | .002 | 9.831583247 | 61.13782921 |
| | 3-Diploid | 6.114177857 | 5.425266922 | .792 | -9.21649540 | 21.44485111 |
| | 4-Triploid | -3.01197138 | 5.559244304 | .982 | -18.7212368 | 12.69729402 |
| | 5-Tetraploid | -2.80037322 | 7.176953535 | .995 | -23.0809476 | 17.48020121 |
| 3-Diploid | 1-H.pup | 29.3705284* | 8.404787374 | .008 | 5.620351494 | 53.12070525 |
| | 2-P.est | -6.11417786 | 5.425266922 | .792 | -21.4448511 | 9.216495396 |
| | 4-Triploid | -9.12614924 | 4.373990027 | .241 | -21.4861332 | 3.233834687 |
| | 5-Tetraploid | -8.91455108 | 6.303590531 | .621 | -26.7271837 | 8.898081582 |
| 4-Triploid | 1-H.pup | 38.4966776* | 8.491885943 | .000 | 14.50037833 | 62.49297688 |
| | 2-P.est | 3.011971379 | 5.559244304 | .982 | -12.6972940 | 18.72123678 |
| | 3-Diploid | 9.126149236 | 4.373990027 | .241 | -3.23383469 | 21.48613316 |
| | 5-Tetraploid | .2115981594 | 6.419262391 | 1.000 | -17.9278991 | 18.35109538 |
| 5-Tetraploid | 1-H.pup | 38.2850794* | 9.628893586 | .002 | 11.07583362 | 65.49432528 |
| | 2-P.est | 2.800373219 | 7.176953535 | .995 | -17.4802012 | 23.08094765 |
| | 3-Diploid | 8.914551076 | 6.303590531 | .621 | -8.89808158 | 26.72718374 |
| | 4-Triploid | -.211598159 | 6.419262391 | 1.000 | -18.3510954 | 17.92789906 |

Based on observed means.
The error term is Mean Square(Error) = 185.431.
*. The mean difference is significant at the 0.05 level.

Supplementary Table 2. Overview of sampling scheme and summary statistics. General linear model and Tukey's honest significance test of multiple comparisons. H. pup = *Halopyrgus pupoides*, P.est = *Potamopyrgus estuarinus*, diploid, triploid, and tetraploid = Varying ploidies of *Potamopyrgus antipodarum*. 'Set' refers to the mean of one three-male replicate.

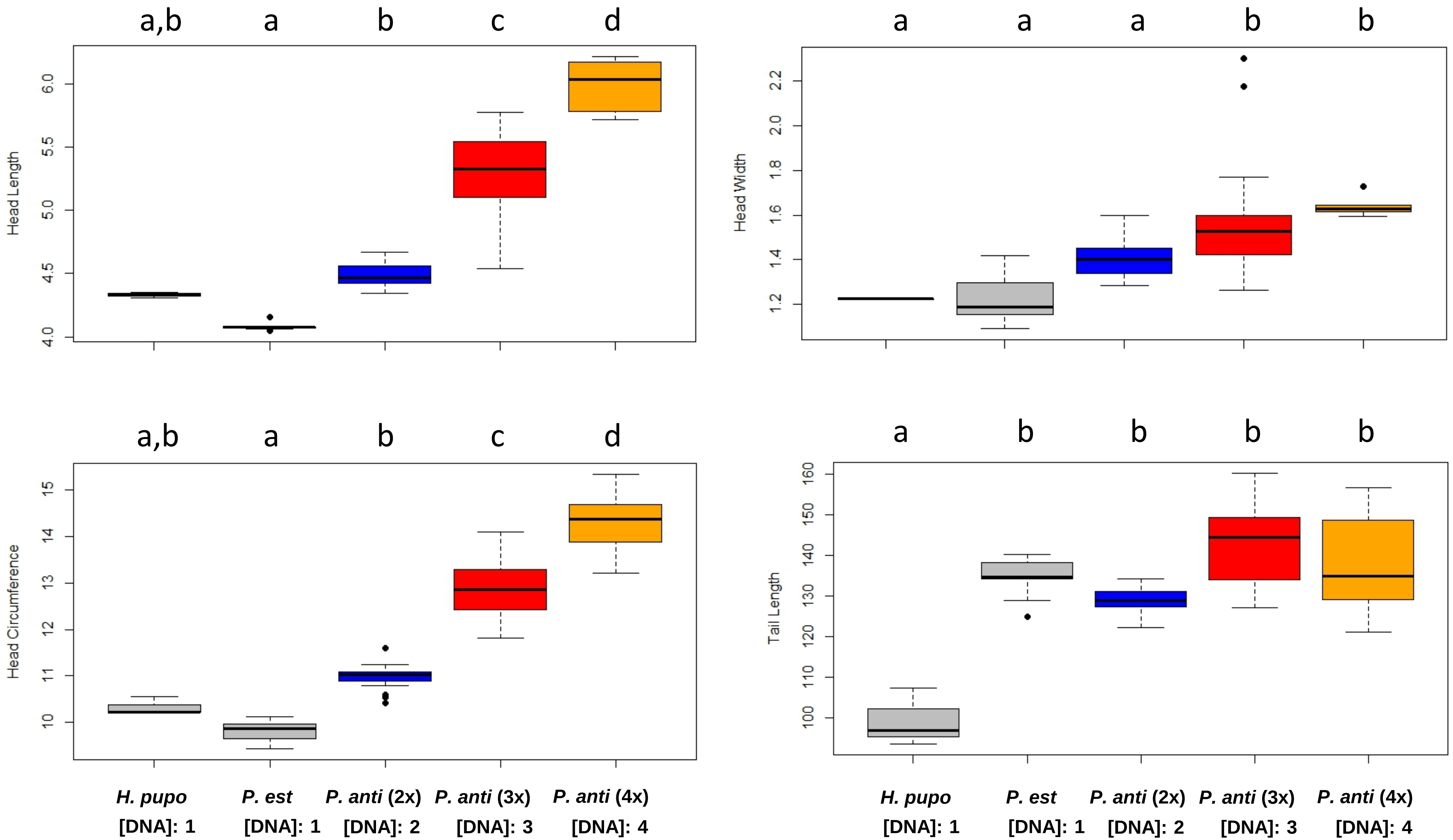


Supplementary Figure 7. Comparisons of sperm traits across nuclear genome DNA contents, ploidies, and reproductive modes. Each data point represents the mean of 12 sperm from one individual male. *H. pupoides* (*H. pupo*) N=3 males, *P. estuarinus* (*P. est*) N=9 males, *P. antipodarum* (*P. anti*) diploid N=21 males, *P. antipodarum* triploid N=18 males, *P. antipodarum* tetraploid N=6 males. We used the independent samples t-test and Tukey post hoc test of multiple comparisons to assess whether the samples were analytically distinguishable. Different lowercase letters at the top of each panel represent statistically different means (Thick black line represents the median, the black box represents first and third inter-quartile range (IQR), whiskers represent Q-1.5 IQR and Q+1.5 IQR, respectively, and black dots represent outliers (> 1.5 times the IQR). “DNA” = relative nuclear DNA content for each category. All measurements are in µm.

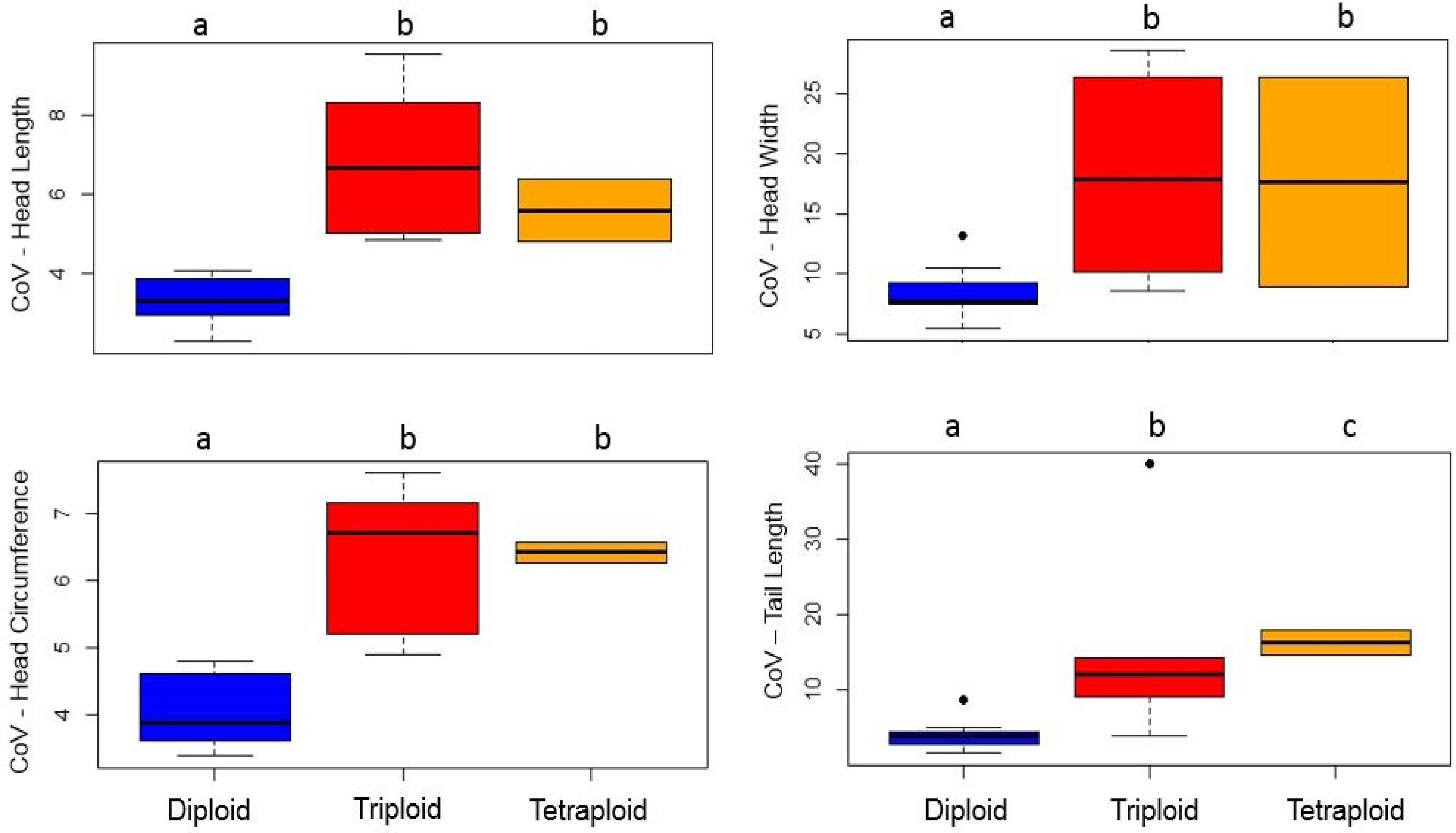


| | | Mean (µm) | SD |
|---|---|---|---|
| HL | Diploid | 4.48 | 0.09 |
| | Triploid | 5.29 | 0.32 |
| | Tetraploid | 5.99 | 0.22 |
| HW | Diploid | 1.41 | 0.08 |
| | Triploid | 1.63 | 0.33 |
| | Tetraploid | 1.73 | 0.21 |
| HC | Diploid | 10.98 | 0.26 |
| | Triploid | 12.86 | 0.58 |
| | Tetraploid | 14.31 | 0.74 |
| TL | Diploid | 128.6 | 3.34 |
| | Triploid | 137.73 | 22.05 |
| | Tetraploid | 137.52 | 12.99 |

Supplementary Figure 8. The effect of sex and ploidy on sperm trait coefficient of variation of each male. *P. antipodarum* diploid N=252, *P. antipodarum* triploid N=216, *P. antipodarum* tetraploid N=72. Different lowercase letters at the top of each panel represent statistically different means ($p < 0.05$). Thick black line represents the median, the black box represents first and third inter-quartile range (IQR), whiskers represent Q-1.5 IQR and Q+1.5 IQR, respectively, and black dots represent outliers (> 1.5 times the IQR).

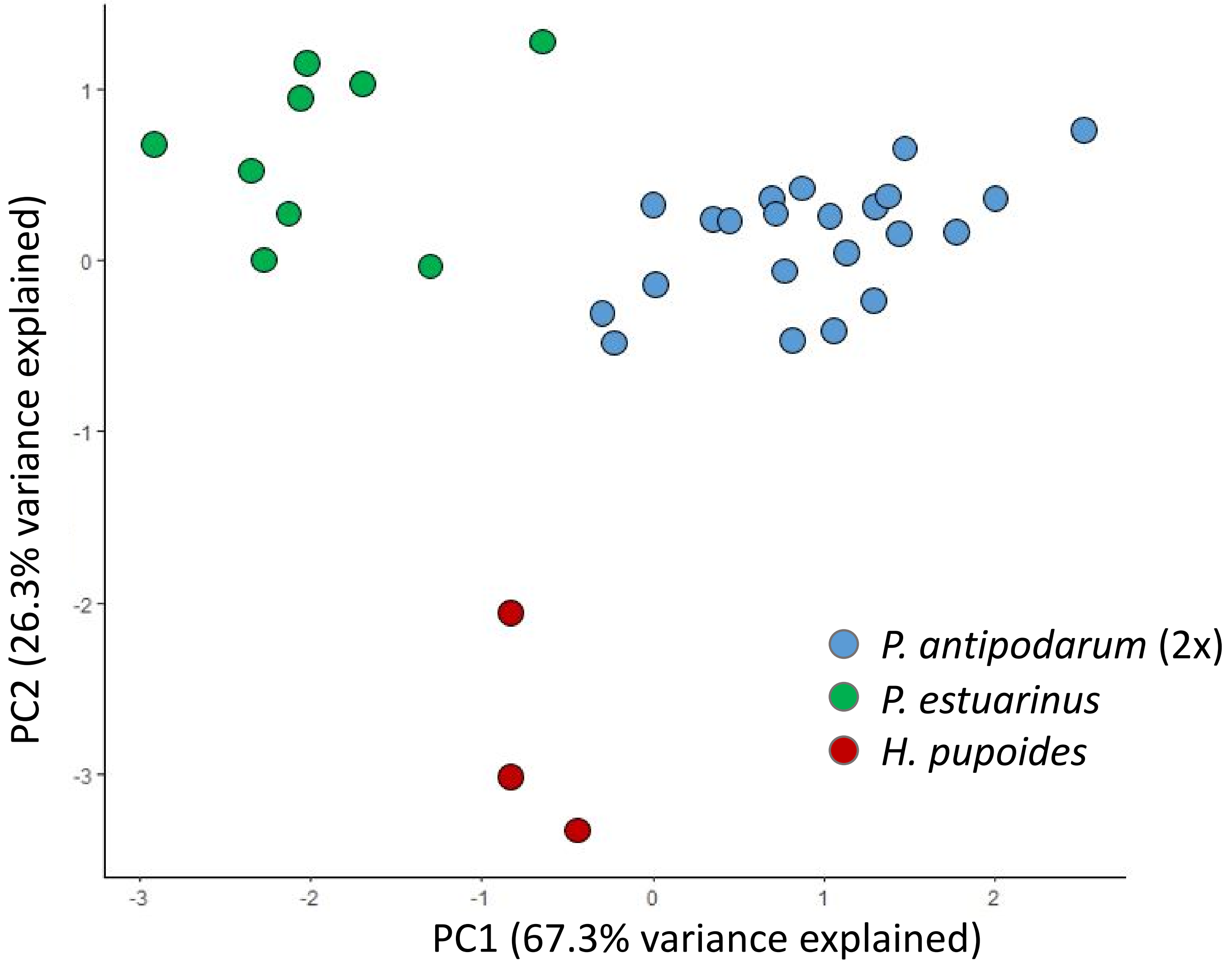


Supplementary Figure 9. Plot of first and second principal components (R 3.4.1) of the mean of transformed sperm traits (head length, head width, head circumference, tail length) for diploid *P. antipodarum* and for all *P. estuarinus* and *H. pupoides*, which are always diploid.

Supplementary Figure 10. Magnitude of difference between sperm traits of particular snail groups as a function of differences in nuclear DNA content between those groups. We generated these "difference" values by dividing 1000 randomly selected (with replacement) sperm measurements from each group by 1000 randomly selected (with replacement) sperm measurements from the other group for each trait. As a control, we also performed the same sampling and comparison within each snail group ($p$ = 1.000 in all within-group comparisons; data not shown). Red bars indicate the null hypothesis of nuclear DNA content driving morphology differences. A value of one indicates that there is no difference between the two lineages. Independent Samples t-tests were used to assess whether the samples were distinguishable using Bonferroni-adjusted $p$=0.0125. 2x=diploid *P. antipodarum,* 3x= triploid *P. antipodarum,* 4x= tetraploid *P. antipodarum, P*est = *P. estuarinus* (~60% the DNA content of diploid *P. antipodarum*). Shared lowercase letters above each panel indicates that comparisons were not statistically distinguishable ($p$ > 0.0125); different letters indicate that the samples differed at the $p$ < 0.0125 level. Even when all four groups were statistically different from one another (top left, bottom left), the differences were not in the rank-order predicted if nuclear genome content values were the main driver. These comparison outcomes rule out nuclear genome content as the sole or main explanation of the differences in sperm traits that we observed across our snail groups.

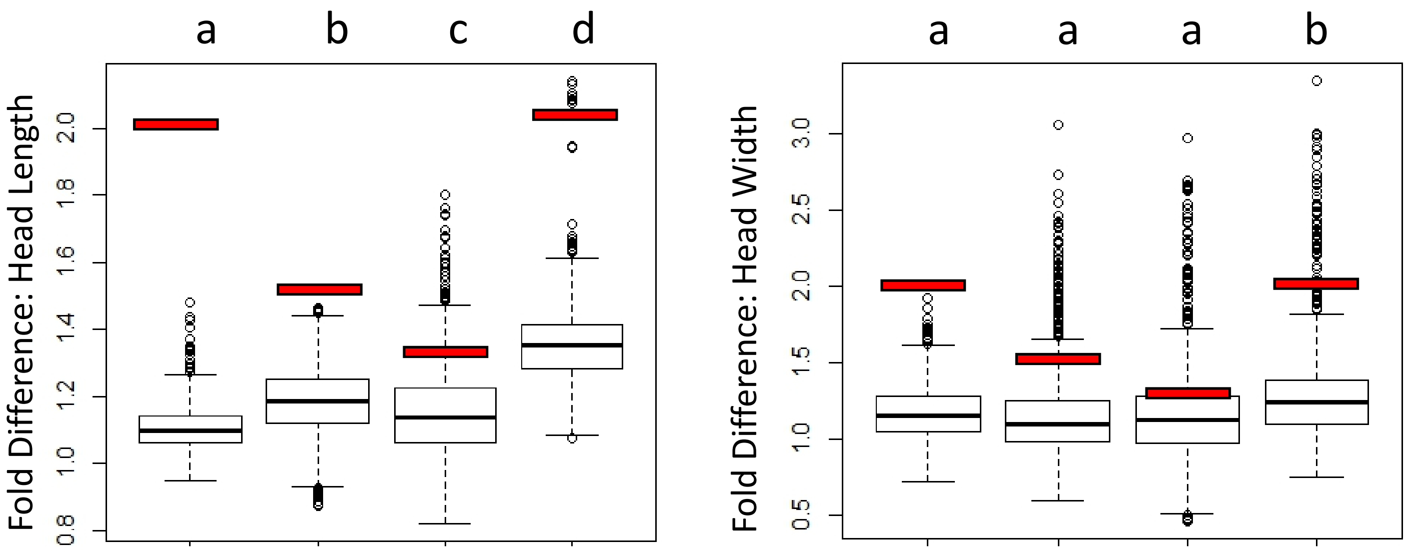


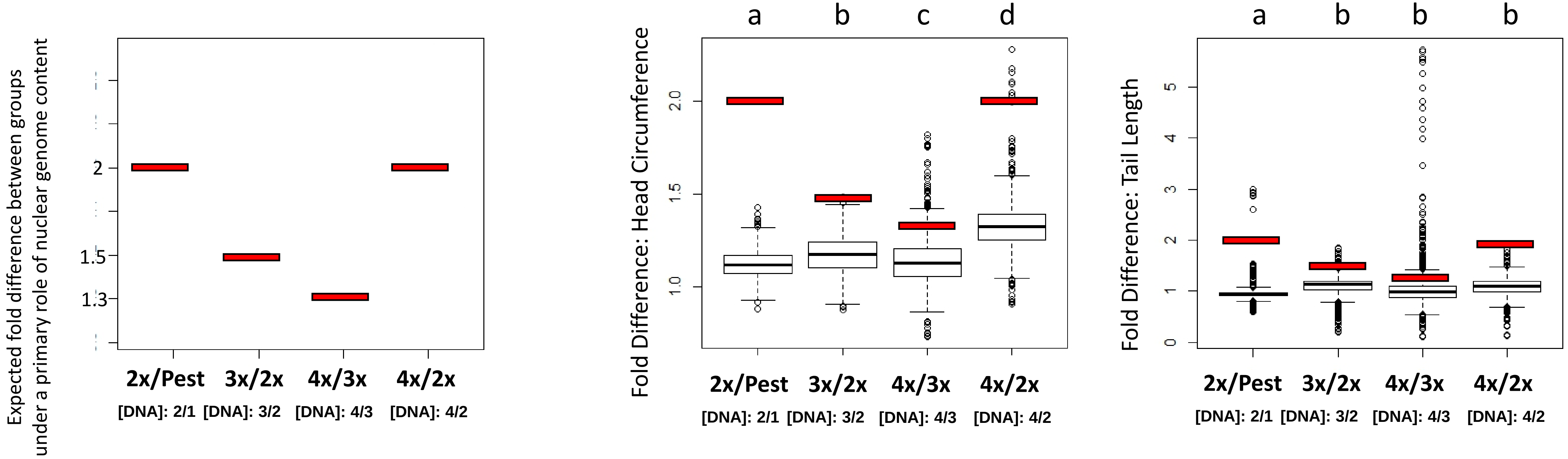
